# Enhanced fitness relates to reduced cerebrovascular reactivity and perfusion in a sample of very healthy older adults

**DOI:** 10.1101/444208

**Authors:** Brittany Intzandt, Dalia Sabra, Catherine Foster, Laurence Desjardins-Crépeau, Rick Hoge, Christopher J Steele, Louis Bherer, Claudine J Gauthier

## Abstract

Aging is accompanied by decreased grey matter volume (GMV), cerebral blood flow (CBF), and cerebrovascular reactivity (CVR), where the latter tends to decline the earliest in aging. Enhanced fitness in aging has been related to preservation of GMV and CBF, and in some cases CVR, although there are contradictory relationships reported between CVR and fitness. Therefore, to gain a better understanding of the complex interplay with fitness and GMV, CBF and CVR, it is necessary to study them concurrently. Here, we aimed to disentangle the interactions between these outcomes in healthy older adults. MRI acquisitions collected anatomical, CBF and CVR information in all participants, as well as VO2,max. Results revealed decreased CVR was associated with increased fitness throughout large areas of the cerebral cortex. Within these regions it was found that lower fitness was associated with higher CBF and a slower hemodynamic response to hypercapnia. Overall, results indicate that the relationship between age, cerebral health and cerebral hemodynamics are complex. Future studies should collect other physiological outcomes in parallel with quantitative imaging, such as measures of carbon dioxide sensitivity and autoregulation, to further understand the intricacy of the effects fitness has on the aging brain, and how this may bias quantitative measures of cerebral health.

## Introduction

During aging, the vascular system goes through a cascade of events that negatively affects the integrity of the cerebrovascular system, leading to decreased perfusion. Continuous and optimal blood flow is thought to be necessary for structural integrity and normal neuronal activity ^1^, so that decreased perfusion likely contributes to the grey matter atrophy^2^ and impaired cognition^3^ observed in aging. It may be possible to reduce these deficits, and in some instances, reverse them due to plasticity. Plasticity refers to the capacity of the brain to change its function, hemodynamics and microstructure in response to experience or physiological changes^4,5^. In aging, there is some indication that physical activity may be capable of inducing beneficial plastic changes^6,7^. Notably, aerobic exercise (i.e. fitness) has become a subject of particular interest for maintaining and even enhancing cognition and brain integrity^7–9^. It is likely that these effects are mediated by changes in cerebral vascular health given the well-established positive influence of exercise on the cardiovascular system in aging^10,11^. Specifically, it has been demonstrated that more highly fit individuals have enhanced endothelial function^12^ and reduced arterial stiffness ^10,13^, both of which are impaired in aging ^14–16^.

Given this positive relationship between the vascular system and exercise, there is an increasing body of work investigating the relationship amongst aging, fitness, structural integrity and cerebral hemodynamics. Magnetic Resonance Imaging (MRI) is a method of choice to study these relationships as it is a versatile technique which can be used to quantitatively measure several of these parameters, including grey matter, cerebral blood flow (CBF) and cerebrovascular reactivity (CVR). In general, grey matter volume (GMV) and CBF are consistently positively associated with fitness in cross-sectional^17–19^ and longitudinal studies^20–24^. However, many of the existing studies showing this beneficial plasticity effect have used GMV as a marker of structural integrity. This is problematic for a physiologically-specific interpretation of these effects since GMV has been shown to be mainly qualitative and physiologically non-specific^25,26^

However, more physiologically specific approaches have involved looking at the relationship between CBF and fitness. In cross-sectional studies, it has been demonstrated that there is a positive relationship between fitness and CBF^18,19,27^. Earlier work by Chapman et al., 2013^21^, found that individuals in a 12-week aerobic training program demonstrated significant improvements to CBF compared to the passive control group. Yet, in a later study Chapman et al. (2016)^28^ found CBF to be unchanged after the same aerobic training program. While this could be due to an insufficient exercise dose, it is possible that CBF is not a sensitive enough or consistent marker of cerebrovascular health in isolation. This could be both due to its relatively limited signal-to-noise ratio^29^ and the fact that homeostasis seeks to maintain CBF to ensure adequate perfusion to maintain oxygen and glucose delivery^30^. In fact, there are indications that while CBF does steadily decrease across the lifespan^31^, more dynamic aspects of hemodynamics, such as cerebrovascular reactivity (CVR) may change earlier than CBF in aging ^32,33^.

CVR is measured as the hemodynamic response (in terms of CBF, increased flow velocity, or blood-oxygen-level dependent (BOLD) change) to a vasodilatory challenge, typically inhalation of CO_2_. CVR is hypothesized to represent the health of the cerebral vasculature^34^ as it could be treated as a vascular vasodilatory capacity biomarker, if it is assumed that CO_2_-related vasodilation signaling is consistent across age and disease groups. If CVR is taken to be a marker of vascular health, it can be posited that those with higher fitness levels would have greater CVR, as their vascular system would be more compliant and have an increased ability to respond to a vasodilatory stimulus. Yet, the literature has found conflicting results, where some have observed increased CVR in relation to higher fitness levels^35–38^, others have found decreased CVR is related to increased fitness ^19^, while others have found no difference ^39–41^ in aging. It is unclear however if this is due to differences in measurement method, spatial localization of the measurement or an interesting physiological interplay between multiple hemodynamic aspects of brain health.

In summary, there are an assortment of negative consequences that can occur due to an aging vascular system, causing deterioration to the cerebral structure and hemodynamics. Importantly though, some studies do seem to indicate that exercise is capable of mitigating some of these adverse complications due to aging. Yet, the fitness literature does suggest that the effects of exercise on brain hemodynamics may be complex, so a more comprehensive imaging approach is necessary to understand the interplay between the effects of aging and fitness on cerebral hemodynamics. Here, we explore the relationship between aging, fitness, cerebral hemodynamics and GMV using a cross-sectional dataset employing a comprehensive imaging approach in healthy younger and older adults with varying levels of fitness.

## Methods

### Participants

A total of 28 young adults (7 females, mean age 24 ± 3 years) and 50 older adults (37 females, mean age 63 ± 5 years) participated in this study and completed a magnetic resonance imaging (MRI) session. Participants were recruited through a previously acquired databases of participants at the Centre de recherche de l’Institut universitaire de gériatrie de Montréal and from Laboratoire D’Etude de la Santé cognitive des Ainés.

Inclusion criteria for participation included being in the age range of 18 to 40 years for young adults and 55 to 75 years for older adults; approval by a geriatrician to participate (older adults), non-smoker (and had for at least five years), no evidence of cognitive impairment as determined through cognitive tests conducted by a neuropsychologist, and completion of the maximal oxygen uptake test (VO_2_max). Exclusion criteria included individuals taking prescription medication known to be vasoactive (e.g. hypertension, statins, thyroid disease, etc.), presence of cardiac disease, hypertensives (including use of anti-hypertensive medication), neurological or psychiatric illnesses, diabetes, asthma, thyroid disorders, smoking within the last 5 years, or drank excessively (more than two drinks per day).

All procedures were approved by Comité mixte d’éthique de la recherche du Regroupement Neuroimagerie/Québec and all participants provided written informed consent.

### MRI acquisition

All acquisitions were completed on a Siemens TIM Trio 3T MRI system (Siemens Medical Solutions, Erlangen, Germany). A 32-channel vendor-supplied head coil was used for all brain acquisitions. An anatomical 1mm^3^ MPRAGE acquisition (TR/TE/flip angle = 300ms/3ms/90°, 256×240 matrix) was acquired for the registration process from native to standard space, and to measure grey matter volume. A fluid attenuation inversion recovery (FLAIR) acquisition with parameters: TR/TE/flip angle 9000 ms/107 ms/120° with echo train length of 15, an inversion time of 2500 ms, 512 x 512 matrix for an in-plane resolution of 0.43 x 0.43 mm and 25 slices of 4.8 mm was used to estimate the presence and severity of white-matter lesions. Further, a pseudo-continuous arterial spin labeling (pCASL) acquisition [25] was acquired, providing simultaneous BOLD contrast using dual-echo readouts (TR/TE1/TE2/flip angle = 2000ms/10ms/30ms/90°) with 4×4×7mm voxels, 64×64 matrix and 11 slices, post-label delay = 900ms, tag duration=1.5s, and a 100mm gap during a hypercapnia challenge (5mmHg end-tidal CO2 change, iso-oxic during two, 2min blocks) were collected.

### Hypercapnia

The hypercapnic manipulation used here has been described previously ^32,42^, however briefly, it was completed with a computer-controlled gas system with a consecutive gas delivery circuit (Respiract™, Thornhill Research Inc., Toronto Canada). The Respiract™ Gen3 provided the ability to independently control the end-tidal partial pressure of oxygen (PO_2_) and carbon dioxide (PCO_2_) by using a feed-forward system based on participants’ baseline or predicted O_2_ consumption and CO_2_ production^43^. End tidal O_2_ was directed to be 100mmHg throughout the manipulation, while CO_2_ had a target of 45mmHg during the hypercapnia blocks and 40 mmHg during normocapnia. Participants breathed through a soft plastic mask (Tegaderm 3M Healthcare, St. Paul MN) that was kept on their face with adhesive tape to ensure that no leaks were present. Participants completed the breathing manipulation once prior to being in the scanner, and once during the MRI session

### VO_2_max

Participants completed a maximal oxygen consumption test (VO_2_max) to approximate their cardiovascular fitness, where a greater amount of oxygen consumed indicates enhanced cardiovascular fitness. The test was completed on a stationary cycle ergometer and were monitored by an electrocardiogram for the duration of the test, which was under medical supervision to ensure participant safety. Initial workload was set based on the body weight of the individual (1 watt (W) / kg) and then increased incrementally by 15 W every minute until voluntary exhaustion. Oxygen uptake was determined using an automated system that averaged in 30-second increments (Moxus, AEI Technologies, Naperville, IL). The highest oxygen uptake over a 30 second period during the test was considered as the maximal amount of oxygen consumed (ml/kg/min).

### Neuropsychological outcomes

A registered neuropsychologist completed the Mini Mental Status Examination which is a global cognitive screening tool for dementia, participants with scores less than 26 (out of 30) were excluded ^44^.

## Data Analysis

### BOLD CVR Data

Preprocessing of the BOLD CVR have been described previously ^42^. Briefly, the BOLD signal was extracted from the second echo series with a linear surround addition. The fractional change within the BOLD during hypercapnia was derived by fitting a general linear model to the BOLD signal and divided by the estimated effect size by the estimated constant term. Glover’s parameters (1999)^45^ were used for the shape of the hemodynamic response function when fitting the models using a single-gamma hemodynamic response function, and included linear, quadratic, and third order polynomials representing baseline signal and drifts.

### Resting CBF

Resting CBF was analyzed using the first echo of the whole pCASL data time series. Given that individual M0 maps were not collected, the average of the control images was used for each participant with modeling of the T1-recovery to obtain the fully recovered magnetization using AFNI, FSL and in-house scripts. Further, cerebrospinal fluid (CSF) masks were created for each older adult participant to use CSF M0 in for CBF quantification. To do this, 10 voxels were chosen in the same axial slice for all older participants where the lateral ventricles were clearly located, except for four participants where a more superior, or inferior slice was required in order to clearly identify the ventricles from the pCASL scans. All individual masks were visually inspected to ensure they were located in the ventricles. For the younger adults, one participant was chosen at random and the same method was used where 10 voxels were chosen in an axial slice where the lateral ventricles were clearly located. A single CSF mask was created for all younger participants as this mask was found to be located in the ventricles in all participants upon visual inspection. FSL’s BASIL^46^ toolkit was then utilized, employing the standard parameters in BASIL toolkit as follows: labelling: cASL/pcASL; bolus duration: constant (1.5s), post label delay: 0.9 s; calibration image: average of the control images; reference tissue type: CSF; mask: CSF mask for each participant; csf T1: 4.3s; TE: 10 ms; T2: 750 ms; blood T2: 150 ms; arterial transit time: 1.3 s, T1: 1.3s, T1 blood: 1.65s, inversion efficiency: 0.85.

### CBF CVR

CBF CVR was calculated in almost the same way as BOLD CVR where the CBF signal was isolated from the first series of echoes using linear surround subtraction^47^. Fractional changes in the CBF signal during hypercapnia was then calculated in the exact same way as was done for the BOLD CVR.

### Structural Outcomes

The volume of white matter hyperintensities for the brain was done in a semi-automatic way as described in^42^. Briefly, a single trained rater, who was blinded to clinical information, visually identified on the fluid attenuation inversion recovery images white matter hyperintensities, which were then delineated using Jim image analysis package, version 6.0 (Xinapse Systems Ltd, Northants, UK).

From the above analyses individual BOLD CVR, resting CBF and CBF CVR were produced for each participant. Co-registration of these maps from native to T1 space was performed using ANTS^48^, and Cat12^49,50^ was used to register from T1 to standard space and for smoothing the structural data with Gaussian filter of 8 mm. An average grey matter mask from each age group was also created in order to analyze all results in grey matter only.

### Aortic Exam

As described in ^45^, during the MRI session a thoracic aortic exam was also acquired using simultaneous brachial pressure recording (Model 53,000, Welch Allyn, Skaneateles Falls, NY USA). Black blood turbo spin echo sagittal oblique images were acquired to visualize aortic arch (TR/TE/flip angle: 700ms/6.5 ms/180°, 1.4 x 1.4 mm in-plane resolution, 2 slices at 7.0 mm), where a perpendicular plane to the ascending and descending aorta was defined from these images at the level of the pulmonary artery. Here, a cine phase-contrast velocity encoded series was collected (TR/TE/flip angle: 28.6ms/1.99ms/30°, 1.5 x 1.5 mm in-plane, one slice at 5.5 mm) during 60 cardiac cycles in three segments, with velocity encoding of 180cm/s in each plane. A series of cine FLASH images were also acquired in the same plane (TR/TE/flop angle: 59ms, 3.44ms, 15°; 1.2 x 1.2mm in-plane resolution, one slice at 6mm) over 60 cardiac cycles in 8 segments.

### Pulse Wave Velocity Data

The aortic data was analyzed using the ARTFUN software^51^, where pulse wave velocity (PWV) was computed from the aortic between the ascending and descending aorta during the cine phase contrast images. PWV was further calculated as completed in ^42^. This data was included again in this paper as a covariate, as we wanted to ensure that arterial stiffness was accounted for so that any relationships that might be present between fitness and the hemodynamic brain outcomes were not due to differences in arterial stiffness among the older adults

### Voxel-wise analyses

Using FSL’s toolbox Randomise^52^; permutation based threshold-free cluster enhancement (TFCE) was employed for analysis in grey matter. Specifically, voxel-wise general linear models were used to identify if relationships existed between VO_2_max and BOLD CVR, resting CBF, or CBF CVR data for young and older adults. Age and sex were included for both young and older adults as covariates. For the older adults we also included a multivariate risk scoring with the Framingham cardiovascular risk factors, as completed by D’Agostino et al. 2008^53^, to estimate general cardiovascular risk and future cardiovascular risk as a confound. Volume of white matter hyperintensities and pulse wave velocity were also used as potential confounds in the older adults.

### Regions of Interest

Regions with a significant relationship between GMV, BOLD CVR, resting CBF or CBF CVR and VO_2_max from Randomise were extracted and binarized to be used as regions of interest (ROI) for further analysis. Specifically, these ROI’s would then be used to further investigate if VO_2_max and GMV, BOLD CVR, resting CBF, or CBF CVR were related to each other in these regions, to further attempt to disentangle the physiological relationship amongst these factors in aging and fitness. This ROI was then multiplied by each individual’s grey matter probability to create individual ROI masks to correct for possible grey matter atrophy. The values from each participant for this individual ROI were then extracted for all maps using weighted average with FSLmeants.

Finally, a finite impulse response (FIR) analysis was completed as reported in ^32^. Briefly, the average time course for BOLD during hypercapnia was determined during the FIR, where the temporal response was measured 15 seconds prior to and after the end of each hypercapnic block in order to identify the zero crossing, the plateau and the linear fit for the values between these two points. These FIR analyses were completed on the significant ROI, where the ROI was applied to each TR, and the value of the BOLD response from the FIR was extracted for all TR’s. This was completed for both young and older adults. The older adult group was further divided using a median split for the fitness data, to allow visualization of the relationship between response shape and fitness.

## Statistical Analysis

Statistical analysis on behavioural data was completed using SPSS 20.0 software (IBM, Armonk, New York, USA). Specifically, to identify if relationships were present between VO_2_max and demographic data, using correlational analyses. A partial correlation was used to identify if there were relationships between VO_2_max and values extracted from significant ROI’s, with only age entered as a covariate into the statistical model. The other above covariates were entered into the linear model in Randomise, however when statistics were completed on the demographic data between higher fit and lower fit older adults, it was found that the only variables that were statistically significant between the two were VO_2_max and age. Thus, as the other covariates (i.e. Framingham Risk Factor Score, log white matter hyperintensities volume, etc.) were already accounted for in the voxel-wise analysis and as they were not statistically different between the older adults and fitness level it was determined that it was not necessary to add these covariates to the model as they should not influence the model.

A one-way ANOVA was utilized with the FIR data to identify if differences were present between young adults, higher fit older adults and lower fit older adults, including the slope of the linear regression between the zero crossing and first plateau in BOLD response. Older adults were split into high fit and low fit using a median split based on VO_2_max. Statistical significance was set to p ≤0.05 for all outcomes and Tukey’s post-hoc analysis was used where applicable.

## Results

### Younger versus older adults

A total of 50 older adults (33F, 17 M) participated in this study. Participant demographics are listed in Table 1.

**Table 1:** Participant Demographics. It was found that younger adults had a significantly higher GMV (p=4.43×10^-15^), BOLD CVR (p=1.4×10^-4^) and resting CBF (p=0.015) in whole GM than older adults. There were no differences between these age groups for CBF CVR in whole GM (p=0.315) (*Table 1*)

| **Demographic** | **Young Adults (n=26)** | **Older Adults (n=50)** |
| --- | --- | --- |
| *Sex (M/F)* | 19/7 * | 17/33 * |
| *Age (years)* | 23.7 (2.9) * | 63.4 (4.9) * |
| *Education (years)* | 16.7 (1.4) | 16.4 (3.6) |
| *VO<sub>2</sub>max (ml/kg/min)* | 42.7 (7.6) * | 29.1 (7.0) * |
| *MMSE (out of 30)* | -- | 28.8 (0.9) |
| *Framingham Risk Factor Score* | -- | 8.8 (2.6) |
| *Log WMH volume* | -- | 0.367 (0.162) |
| *Grey Matter Volume (mm<sup>3</sup>)* | 0.551 (0.038) * | 0.466 (0.034) * |
| *BOLD CVR (%change/mmHg CO<sub>2</sub>)* | 0.261 (0.094) * | 0.176 (0.041) * |
| *Resting CBF (ml/100g/min)* | 48.6 (10.7)* | 42.4 (9.9)* |
| *CBF CVR (ml/100g/min/mmHg CO<sub>2</sub>)* | 5.13 (1.22) | 4.63 (2.35) |

### Fitness and hemodynamics in older adults

The mean values in all GM for each participant are shown in Figure 1 demonstrating their relationship with VO_2_max for both young and older adults. ROI analyses of all GM within the younger adults did not reveal any significant relationships between VO_2_max and GMV or between VO_2_max and any of the hemodynamic outcomes (p >0.05). In older adults, it was identified that GMV was positively related to VO_2_max (r=0.320; p=0.025) and BOLD CVR had a negative association (r=-0.392; p=0.005). No relationship was found between VO_2_max and resting CBF or CBF CVR (p>0.05).

**Figure 1:**
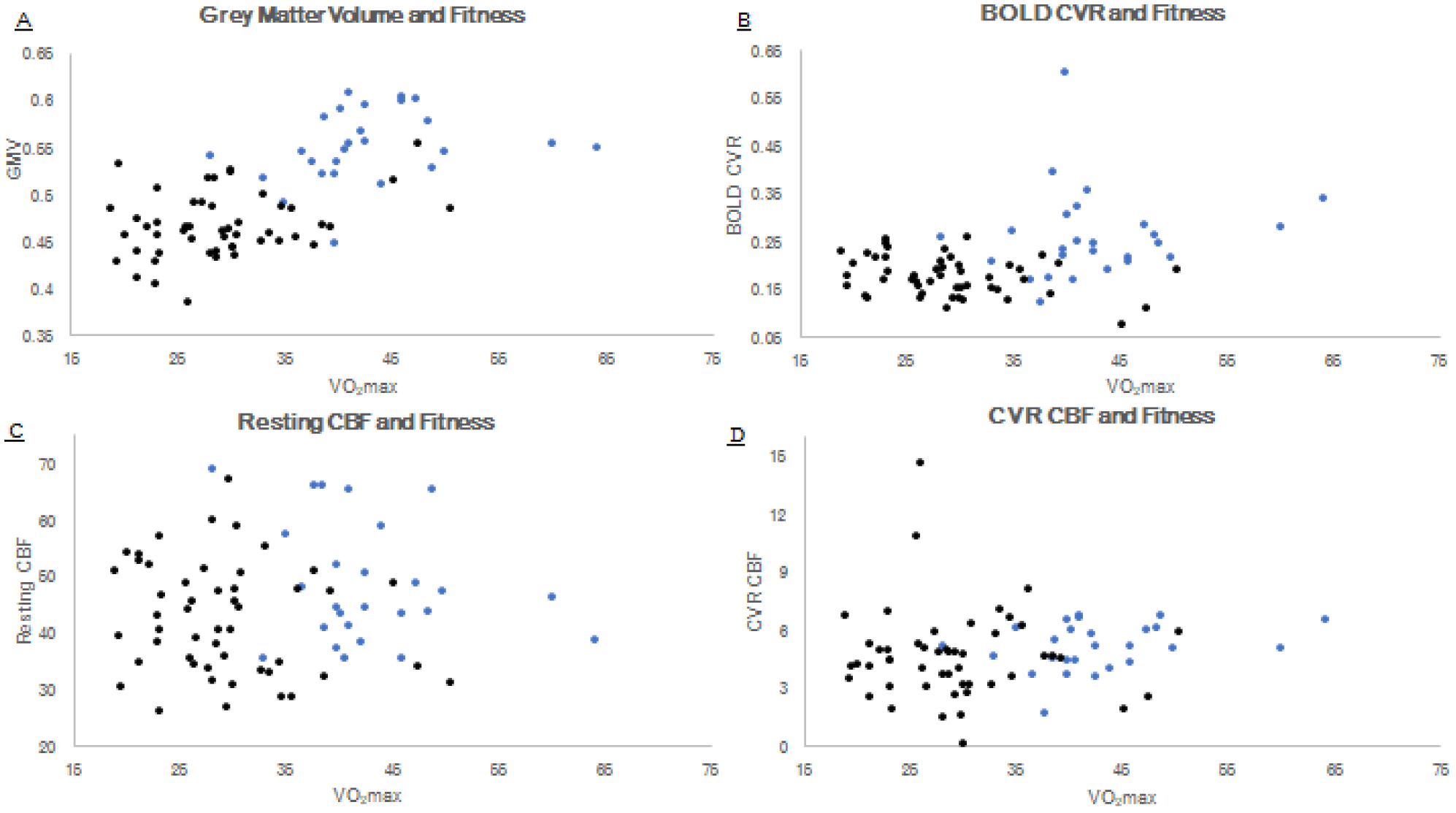
Relationships for between structural and hemodynamic outcomes with VO2max (ml/kg/min) for young adults (black dots) and older adults (blue dots). A: grey matter volume (mm3); B: BOLD CVR (%BOLD/mmHg CO2); C: resting CBF (ml/100g/min) and; D: CVR CBF (ml/100g/min/mmHg CO2)

To understand the regional relationships between fitness and these structural and hemodynamic parameters, a voxel-wise analyses were performed. Voxel-wise analysis demonstrated a significant negative relationship between BOLD CVR and VO_2_max in the regions demonstrated in Figure 2. No regions with a significant relationship with VO_2_max were found for GMV, resting CBF or CBF CVR (p>0.05).

**Figure 2:**
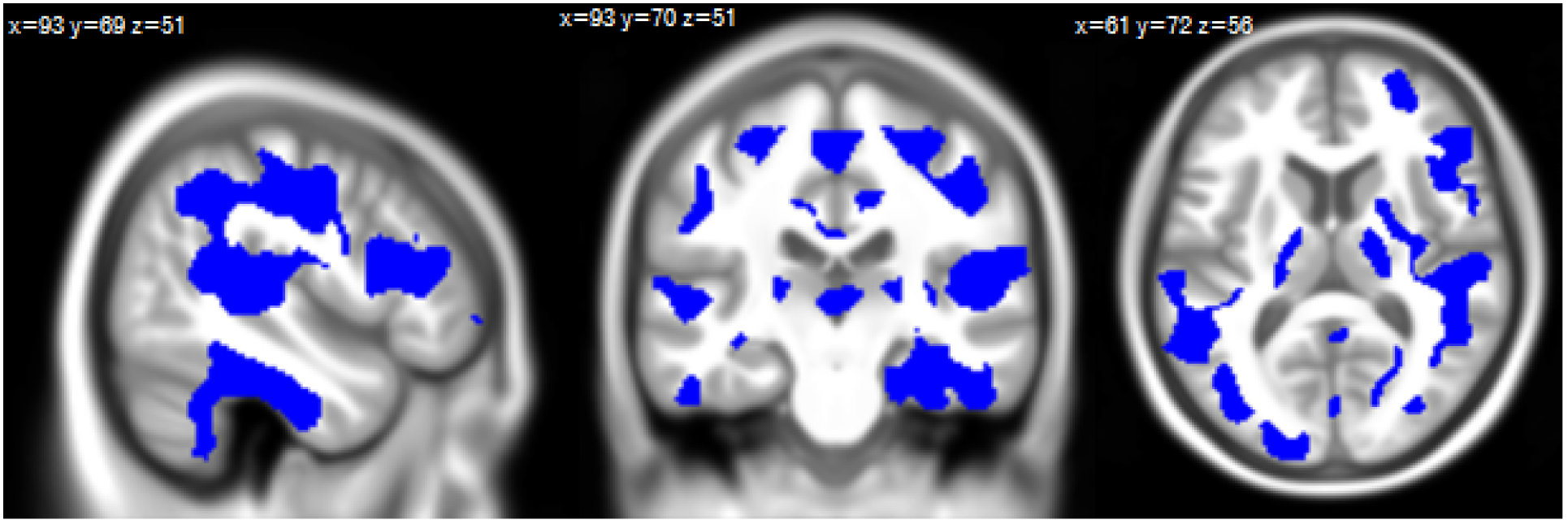
Areas of the brain where there is a negative association between VO2max and CVR, indicating that those with higher fitness, in these areas, have significantly reduced BOLD CVR compared to those with lower fitness (p<0.05)

The regions that were found to be significantly related for BOLD CVR and VO_2_max were then used to create an ROI. This ROI was used to extract mean GMV, resting CBF, and CBF CVR within these regions to examine the relationship between VO_2_max and the other hemodynamic parameters. A significant negative relationship was identified between VO_2_max and resting CBF (r=-0.313, p=0.029). The correlation between VO_2_max and CBF CVR flow approached significance (r=-0.280, p=0.052). All relationships between fitness and structure or hemodynamics are shown in Figure 3.

**Figure 3:**
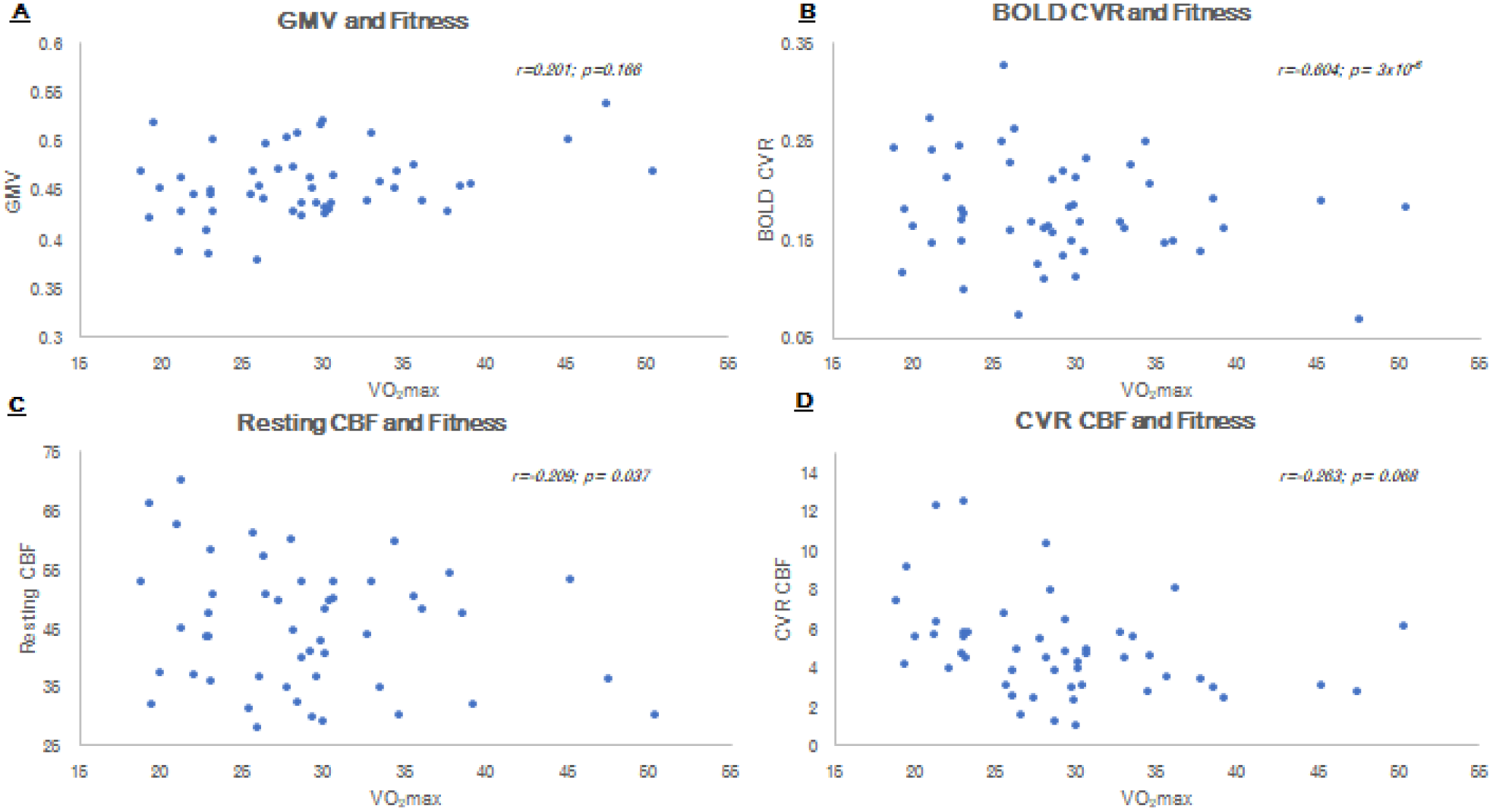
Relationships from the CVR VO2max ROI in: A GMV; B: BOLD CVR; C: Resting CBF; and D: CBF CVR with fitness extracted from the for all older adults.

Finally, to identify potential relationships between fitness and BOLD response dynamics, a Finite Impulse Response (FIR) analysis was completed in the significant CVR VO_2_ ROI. For visualization purposes, the BOLD time course to hypercapnia in these areas amongst young adults, highly fit older adults and lower fit older adults (identified through a median split approach) are demonstrated in Figure 4a. The relationship between the slope of the upward response and VO_2_max is shown in Figure 4b. A partial correlation was completed to investigate in the older group if there was a relationship between VO_2_max and the slope of the BOLD times to hypercapnia (age was a covariate). This correlation revealed that there was a significant negative correlation between VO_2_max and the slope of the linear regression (r=-0.441; p=0.002) (Figure 4B).

**Figure 4A:**
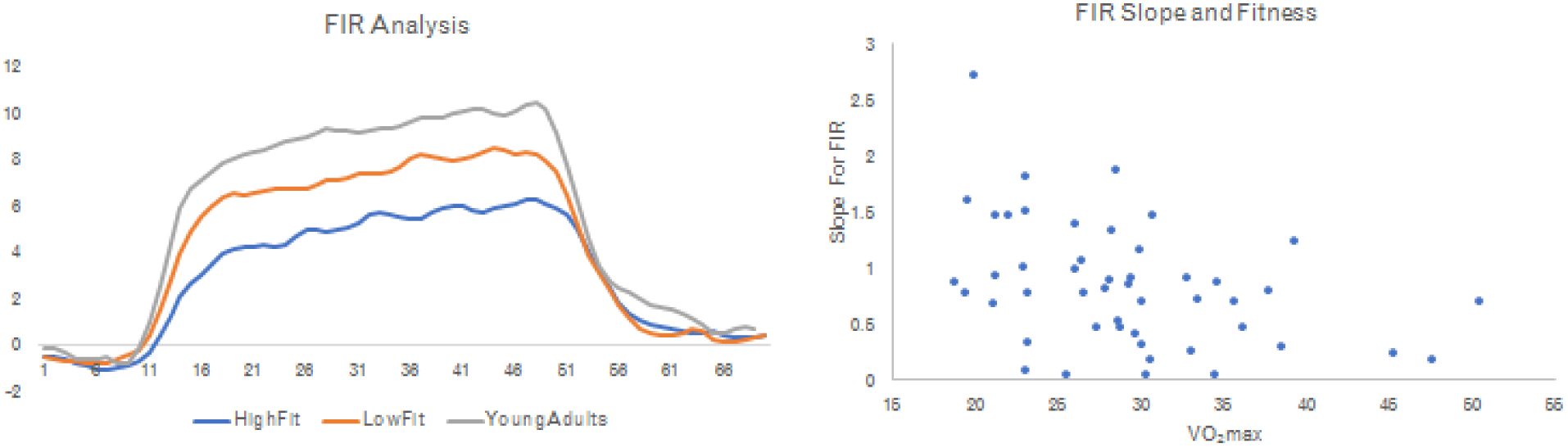
Percent blood oxygen level dependent (BOLD) signal to hypercapnia. Time course showing BOLD response to hypercapnia in CVR VO2max ROI. B: Linear regression between time to pass 0 and time to first plateau and the relationship with fitness in older adults

## Discussion

This study investigated the relationship between fitness, GMV and brain hemodynamics in a highly healthy population of older adults. This group of older adults showed the expected pattern of reduced GMV, BOLD CVR and resting CBF as compared to healthy younger adults. ROI analysis over all GM demonstrated a significant positive relationship between fitness and GMV and a negative relationship between BOLD CVR and fitness. Surprisingly however, regional analyses revealed a significant inverse relationship between BOLD CVR and fitness in a number of GM regions throughout cortex. A more in-depth review of hemodynamics within these regions showed that the relationship between fitness and other hemodynamic parameters also exhibited this inverse relationship. This follow-up analysis revealed no relationship between fitness and GMV, but a significant negative relationship between fitness and resting CBF, and a trend towards a negative relationship with CBF CVR. To determine whether these relationships between fitness and BOLD CVR were confined to response amplitude or if it was also present in response dynamics, we performed an analysis of the time course of the BOLD response to hypercapnia. This time course analysis revealed a slower ramp-up towards a plateau in more highly fit older adults as compared to both less fit older adults and younger adults. Together, these results indicate that the relationship between fitness and hemodynamics in aging is more complex than previously thought and that BOLD CVR in particular may be biased by physiological mechanisms affected by exercise.

The impact of healthy aging on brain structure and function is an active field of research and the age group comparisons performed as part of this study are consistent with these existing results. Older adults were found to have a lower GMV volume as compared to young adults, similar to previous reports of age-related atrophy in GM ^54–56^. Our findings of decreased CVR in older adults compared to young adults is also in line with previous research ^32,33,39,57,58^, as are the findings of decreased resting CBF^32,33^. Some of these results are similar to previously published results by our team in this same group of subjects^32,42^. These age comparison results are presented here despite this previous report both as support and contextualization for the fitness results, and because improvements in slab registration have led to more reliable group results than in previous publications of these results.

The main result from this study was the finding that BOLD-CVR in GM is negatively correlated with fitness in older adults. Voxel-wise analysis revealed large sections of GM including temporal, parietal and frontal regions were responsible for this negative relationship. Given that higher CVR is typically interpreted as being related to better cerebral health^34^, this reverse relationship was counter-intuitive. Interestingly, to date, only one other published study (in addition to our own previous work) has also demonstrated a negative relationship between fitness and BOLD CVR in older adults^19^. In the Thomas et al. study, Master athletes were found to have a significantly reduced BOLD CVR compared to their sedentary counterparts over most of cerebral cortex. While most studies into these effects have identified a positive relationship between CVR and fitness ^35–38^, these studies used transcranial Doppler (TCD) to measure effects.

MRI studies have so far mainly identified a negative relationship, perhaps reflecting the different vascular compartments and properties imaged with both techniques. TCD images flow velocity in major arteries, while the BOLD signal reflects a mixture of blood flow, blood volume and oxidative metabolism arising from the parenchyma and veins. Therefore, it is possible that changes in the venous system or related to the parenchymal vasculature leads to these BOLD-CVR results.

ROI analysis over the entire GM also revealed a positive relationship between fitness and GMV. This is consistent with other studies pointing to an association between fitness and GMV^9^. However, as mentioned previously, GMV should be interpreted with caution as it is qualitative, not physiologically specific and may be biased by differences in blood volume^25,26^.

Notably, the only hemodynamic outcome to be significantly associated with fitness in the voxel-wise analysis was the BOLD CVR. The lack of relationship with other hemodynamic parameters could be attributable to the fact that this is a highly healthy group of older adults. Exclusion criteria were numerous and was comprised of taking medication regularly, suffering from cardiovascular risk factors or chronic diseases. Furthermore, the Framingham scores for this group is low, with the average (8.8) just below the score expected purely because of the average age of the group (9), indicating the overall absence of cardiovascular risk factors within the group. Thus, this group might not be a representative sample of “normal” aging, but perhaps more an example of “healthy” aging.

Given this information, this highlights the fact that CVR may in fact be one of the first hemodynamic properties to decline in aging, indicating it’s greater sensitivity to aging-related changes in the cerebrovasculature than CBF and GMV^2,3^. Furthermore, voxel-wise analyses require stringent thresholding techniques to overcome the problem of large number of multiple comparisons being tested. Therefore, it is possible that more subtle regional relationships in CBF or GMV are not detected using this technique.

To better understand the physiological underpinnings of this negative relationship between fitness and CVR, a more in-depth investigation of the relationship between fitness and other hemodynamic and structural parameters within these regions was performed. No relationship between GMV and fitness was revealed within these areas. Within the clusters relating BOLD CVR and fitness, resting CBF and CBF CVR both also had a significantly negative association with fitness, indicating that lower fit individuals had higher CBF and CBF CVR. These findings are in opposition to those reported by Zimmerman et al^18^., and Tarumi et al.^27^, where higher fitness was associated higher global, frontal and parietal CBF, and higher CBF in the occipitoparietal area in those with higher fitness *and* decreased carotid stiffness. Yet, in the Zimmerman study 39% of their participants were on blood pressure medication, thus it could be that medication impacted these results given the vasoactive nature of these medications. Furthermore, this indicates that their higher fit individuals are more representative of our lower fit participants in this study, indicating a possible non-linear relationship between CBF and cerebrovascular health. The results from Tarumi may indicate that arterial stiffness may impact the relationship between fitness and CBF. However, this would require a large cohort study of varying fitness levels, medication use and arterial stiffness in aging to determine the interplay between all these parameters. Future studies should seek a more inclusive recruitment strategy and larger cohort size to address these issues.

Interestingly, the ROI based on the negative BOLD-CVR and fitness relationship also includes regions such as large sections of the temporal and parietal cortices. This is interesting since temporal and parietal cortices are key regions that demonstrate *increased* blood flow in individuals that are APOE4 carriers^59–61^, but longitudinally have significantly decreased flow compared to those without the allele^61^. This is thought to indicate maladaptive compensation in individuals at an increased risk for Alzheimer’s. Furthermore, studies have identified a similar pattern of hyperperfusion in both mild cognitive impairment and in Alzheimer’s patients as compared to healthy older adults, in areas of the temporal and, in some cases, the parietal cortex and the hippocampus^62,63^. Finally, recent work has demonstrated that Alzheimer’s disease patients have a significant decrease in oxygen extraction fraction compared to healthy controls^64^, supporting previous work suggesting there is a greater reduction in metabolism than flow^65^, suggesting that there could be a decoupling between flow and metabolism^62^.

While our individuals are healthy, our results could still be informed by these observation in Alzheimer’s since it is likely that these changes in perfusion and OEF are in fact present within healthy individuals since they were observed in healthy ApoE4 carriers. Furthermore, as these are relative levels of hyperperfusion always observed in relation to a healthier group, it is possible that these relative differences are also present within a group of overall healthy older adults, with fitness being a means of separating individuals which are healthier than others. Therefore, while we do not mean to imply that the lower fit individuals in our group are more likely to develop dementia, it is possible that these patterns of relative “hyperperfusion” are in fact a very early marker of reduced cerebrovascular and metabolic health also present within the healthy population. Given the greater relationship between BOLD CVR and fitness within these areas, it is also possible that “elevated” BOLD CVR in these regions precedes the appearance of relative “hyperperfusion”. This remains to be tested through a longitudinal study however, as this may also simply be an effect of greater signal-to-noise ratio in BOLD-CVR than in ASL-based measurements of perfusion. Taken together, it is possible that the increased resting CBF and CBF CVR in the temporal, parietal and slight frontal regions in lower fit individuals in this study might indicate that there is likely a continuum of maladaptive compensation occurring between metabolism and CBF even in healthy aging. Future work should concentrate on investigating further this decoupling between cerebral metabolism and flow in healthy older adults with a broader range of fitness levels to attempt to further disentangle this relationship with fitness.

In addition to the relationship between BOLD-CVR and fitness already discussed and visible in Figure 4A as a difference between response amplitude between more highly fit and less highly fit (median split) older adults, we have also identified a relationship between the slope of the upswing of the response to hypercapnia and fitness. The slope of the return to baseline was not however related to fitness. This slower BOLD response to hypercapnia in more highly fit older adults could be indicative of a desensitization to CO_2_. While this slower response could theoretically also be due to pre-dilation^66^, this is unlikely to be the case here as lower resting CBF was found to be linked to higher fitness within these same regions. Further studies including additional measurement of autoregulation and the respiratory response to exercise for example could help untangle the physiological underpinning of this response dynamics.

Overall, there are a few rationales that could explain our results of low BOLD CVR in higher fit individuals. The first was hypothesized by Thomas and colleagues ^19^, that perhaps higher fit individuals have decreased sensitivity to CO_2_ in their cerebral blood vessels due to, likely, a lifetime of increased exposure because of increased aerobic activity. In fact, this is consistent with studies showing that endurance training reduces the ventilatory response at a given workload, indicating a decreased local chemosensitivity^67,68^. Nitric oxide is the primary mechanism to respond to changing pH levels from CO_2_ in an attempt to maintain homeostasis in the brain^69,70^. It is thus possible that higher fit individuals could have decreased levels of nitric oxide in response to hypercapnia, which could explain the reduced blood flow response to hypercapnia. Moreover, it has also been proposed that cerebral inflammation could lead to increased nitric oxide signaling^71^, thus lower fit individuals could have increased nitric oxide due to the presence of inflammation. This would also be consistent with the higher resting CBF observed in lower fit individuals. The presence of inflammation is however unlikely to be the main explanation for our results, given the fact that this cohort is overall very healthy, even in the lower fit group. Lastly, nitric oxide also seems to have an effect on cerebral autoregulation^72^, thus when present, it increases the ability of the cerebral blood vessels to dilate in response to a sudden drop in blood pressure, allowing for sufficient blood to flow. Importantly, exhaustive exercise has been found to impair cerebral autoregulation^73^, and was observed to be attenuated in young master athletes compared to sedentary counterparts ^74^. It is therefore possible, given the interplay between decreased CO_2_ sensitivity, nitric oxide presence and cerebral autoregulation, that in combination, in aging, this may account for the reduced BOLD CVR in higher fit individuals.

The results of this and other studies have shown that quantitative techniques such as MRI measurements of CVR and CBF may be biased by health components typically not taken into account in MRI studies, such as fitness and possibly cerebral autoregulation. This is problematic as it may lead to bias in group comparisons or longitudinal studies. Although it is not possible to test any of these outcomes here, it highlights the need for comprehensive studies that seek to measure all the components of the complex relationship between cerebral hemodynamics and fitness. These studies are necessary to make these techniques truly quantitative and reveal the true changes that occur in aging and disease.

## Limitations

Although we found that increasing fitness is related to decreased BOLD CVR in aging, it is difficult to attempt to interpret our results in comparison to other studies due to the high level of variability BOLD CVR in aging literature. For example, other studies report 0.19% BOLD/mmHg in line with our results^75^, yet others report higher levels at 0.28%^76^ and lower levels at 0.13%^77^. This indicates that there is physiological variability within individuals and between studies, as well as technical variability (i.e. type of scanner, delivery of CO_2_, amount of CO_2_ inhaled), thus more work is necessary to further comprehend inter-individual variability, and as noted previously, more robust study designs with a greater breadth of outcome measures.

A limitation of the use of ASL for measuring CBF is that extensive coverage of the entire brain is typically not possible without advanced parallel imaging techniques. Given that the original aim of this study was more focused on executive functions and the frontal lobes the ability to also capture structural and hemodynamics of the hippocampus was not possible here. Therefore, while the hippocampus is a structure associated with fitness-related changes^9^, we are unable to test these types of associations here. Thus, future work aiming to disentangle the relationships among aging, cognition and fitness should optimize the acquisition to capture both the entire cerebral cortex and the hippocampus.

Another limitation to this study is its cross-sectional design, thereby reducing the ability to draw clear conclusions about the relationships between fitness, aging and brain health. While large longitudinal cohorts exist, none have so far also included measurement of CVR and VO_2_max. This is likely because both these techniques are challenging to implement. However, the inclusion of ASL is becoming more common and future studies could attempt to use these cohorts for studying the relationship between physical activity, CBF and other measures of vascular health. Dedicated longitudinal studies over several years including VO2max, CBF and CVR would however be necessary to truly understand these relationships.

Finally, as discussed previously, it is of utmost importance for future studies to include other physiological outcomes, such as measures of CO_2_ sensitivity, or autoregulation, as well as the role that cerebral blood volume, oxygen extraction fraction, cerebral metabolic rate of oxygen consumption and other hemodynamic outcomes. The results presented here suggest that several physiological mechanisms could be at play and measurement of these other physiological outcomes is crucial to understand the physiological origins of the link between cerebral hemodynamics and fitness. This is not only an important component of our understanding of the impact of fitness on the brain, but is also necessary for measurements of CVR and CBF to be truly interpretable as quantitative representations of brain health. The results presented here suggest that less healthy older adults may have a seemingly preserved CBF and CVR as compared to healthier older adults. Therefore, it is possible that existing studies of aging and disease which use these quantitative techniques have in fact underestimated the impact of aging and disease on cerebral hemodynamics. Future studies of the impact of fitness on different peripheral measures of chemosensitivity and autoregulation for example could be used to determine the best measurements to correct or estimate the bias in individual CVR and CBF, or create normative datasets to estimate this bias.

## Conclusions

Overall, this paper identified that there is a negative relationship between BOLD CVR and fitness in a very healthy older adult sample. Within the ROI’s that demonstrated a significant a significant relationship it was identified that other hemodynamic outcomes also showed negative relationships with fitness. These negative relationships could be the result of changes in CO2 sensitivity, or autoregulation changes for example. These results suggest that quantitative measures of CVR and CBF could be biased by unknown physiological changes in these autoregulatory and chemosensitivity properties, and that studies using these markers in aging and disease may underestimate their effects on cerebral hemodynamics. Thus, to further understand and attempt to disentangle the modulatory effect that fitness has on hemodynamics in aging, more comprehensive studies of physiological outcomes are necessary.

## Acknowledgements

The authors thank Carollyn Hurst and André Cyr for their help with data acquisition, Élie Mousseaux, Alban Redheuil, Muriel Lefort, Frédérique Frouin, and Alain Herment for their help with the aortic protocol and analysis, Cécile Madjar, Mélanie Renaud and Élodie Boudes for their help with logistics, and Fatemeh Razavipoor and Julia Huck for helpful discussions. The authors thank Saïd Mekary for his help with VO2max testing, Ellen Garde, Arnold Skimminge and Pernille Iversen for their help with vascular lesion segmentation. They thank Céline Denicourt for performing the blood draws. They thank Jiongjiong Wang of the Department of Neurology at UCLA who provided the dual-echo pseudo-continuous arterial spin labeling sequence. This work was supported by the Canadian Institutes of Health Research (MOP 84378, Banting and Best Scholarship held by C.J. Gauthier), the Canada Foundation for Innovation (Leaders Opportunity Fund 17380), the Ministère du développement économique, de l’innovation et de l’exportation (PSR-SIIRI-239), the Canadian National Sciences and Engineering Research Council (R0018142, RGPIN 2015-04665), the Heart and Stroke Foundation of Canada (New Investigator Award held by C.J. Gauthier).

